# Ectomycorrhizal fungi are influenced by ecoregion boundaries across Europe

**DOI:** 10.1101/2024.03.06.583687

**Authors:** Guillaume Delhaye, Sietse van der Linde, David Bauman, C. David L. Orme, Laura M. Suz, Martin I. Bidartondo

## Abstract

**Aim:** Ecoregions and the distance decay in community similarity are fundamental concepts in biogeography and conservation biology that are well supported across plants and animals, but not fungi. Here we test the relevance of these concepts for ectomycorrhizal (ECM) fungi in temperate and boreal regions.

**Location:** Europe.

**Time period:** 2008 – 2015.

**Major taxa studied:** Ectomycorrhizal fungi.

**Methods:** We used a large dataset of ∼ 24,000 ectomycorrhizas, assigned to 1,350 operational taxonomic units, collected from 129 forest plots via a standardised protocol. We investigated the relevance of ecoregion delimitations for ECM fungi through complementary methodological approaches based on distance decay models, multivariate analyses, and indicator species analyses. We then evaluated the effects of host tree and climate on the observed biogeographical distributions.

**Results:** Ecoregions predict large-scale ECM fungal biodiversity patterns. This is partly explained by climate differences between ecoregions but independent from host tree distribution. Basidiomycetes in the orders Russulales and Atheliales and producing epigeous fruiting bodies, with potentially short-distance dispersal, show the best agreement with ecoregion boundaries. Host tree distribution and fungal abundance (as opposed to presence/absence only) are important to uncover biogeographical patterns in mycorrhizas.

**Main conclusions:** Ecoregions are useful units to investigate eco-evolutionary processes in mycorrhizal fungal communities and for conservation decision-making that includes fungi.

## Introduction

The study of biogeographic patterns is central to understand the ecological and evolutionary processes that shape ecosystems. However, microbes tend to exhibit less clear or different biogeographic patterns compared to plants and animals (Fierer, 2008; Tedersoo et al., 2022; Vasar et al., 2022). For example, the decay of community similarity over increasing geographic distances (or ‘distance decay’, Soininen et al., 2007) could be less relevant for microbes than for other organisms, and most microbial species could have a near-global distribution, being filtered only by local environmental conditions and biotic interactions (Locey et al., 2020; Meyer et al., 2018). However, blurry biogeographical patterns observed in microorganisms could also be due to differences in taxonomic resolution, spatial scale, sampling effort and/or the sampling of inactive individuals (e.g. spores) in microorganism communities (Meyer et al., 2018). Many large-scale studies rely on different sources of data and sampling protocols, such as environmental DNA (eDNA), fungal fruiting body production and online databases compiling data from multiple sources, leading to variability in species detection and identification mistakes (Maldonado et al., 2015). Reliable large-scale data on symbiotic microbial communities is notoriously difficult to obtain because these organisms are typically hidden within their substrates or hosts (e.g. soil, plants), and they have a large and still mostly unknown diversity (Niskanen et al., 2023). In addition, species abundances and local diversity generally cannot be reliably inferred from databases of species distribution data (e.g. Global Biodiversity Information Facility - GBIF.org) or eDNA (Janowski and Leski, 2023). This leads to ignoring a crucial component of diversity by underestimating the importance of abundant species while overestimating the contribution of rare species (Jost et al., 2011).

The concept of ecoregion (“regions of relative homogeneity with respect to ecological systems involving interrelationships among organisms and their environment”, Omernik, 1995) is an important biogeographical tool for conservation. Ecoregions are strongly predictive of species distributions in macroecology, with studies showing that distribution patterns across plant and animal taxonomic groups are most often linked with global ecoregion boundaries (Smith et al., 2020, 2018). Comparatively, fungi appear to be less defined by ecoregion boundaries, which is coherent with the hypothesis that most fungi are ubiquitous and free of dispersal barriers at large spatial scales (O’Malley, 2008).

Ectomycorrhizal (ECM) fungi represent an interesting case study for biogeographic distribution. They are keystone organisms, playing a pivotal role in ecosystems by contributing to plant establishment, tree nutrition, nutrient cycling and carbon sequestration (Smith & Read, 2008; Hawkins et al., 2023). Compared to unicellular or saprotrophic fungi, ECM fungi are usually large and long lived (Douhan et al., 2011), and display a common trophic mode. Ectomycorrhizal communities display marked fine scale structure at the landscape level due to fine scale variation in soil physicochemical properties, dispersal limitation and biotic processes (Bahram et al., 2015; Bauman et al., 2016). However, ECM communities are also structured at biogeographical scales by environmental factors such as climate and the distribution of suitable hosts (Põlme et al., 2018; Tedersoo et al., 2014a; van der Linde et al., 2018). Interestingly, these large-scale biogeographical patterns can contrast with those observed in plants and animals. For example, ECM diversity peaks in temperate and boreal biomes instead of tropical ones (Soudzilovskaia et al., 2019; Tedersoo et al., 2014a). Finally, speciation associated with phylogenetic niche conservatism can also lead to patterns of community clustering at biogeographical scales in fungi (Tedersoo et al., 2014b).

In this study, we investigate spatial and macroecological patterns of ECM species distributions across Europe using a high-resolution dataset of abundances of ECM fungal operational taxonomic units (OTUs). Using a robust method based on the estimation of differences between parameters of distance-decay curves, coupled with multivariate and indicator species (IndVal) analyses, we 1) tested whether and to what extent ECM fungal community composition is defined by distance decay and ecoregions; 2) explored the role of host species distribution, ECM fungal dispersal traits and phylogeny, and 3) investigated the effect of species abundance on the observed patterns. We expected to find:

i. a negative exponential distance decay of ECM fungal community similarity across Europe, and an ecoregion effect on communities’ similarity comparable to what is observed in other organisms (i.e. ECM communities within ecoregions are more similar than communities between ecoregions) but mediated by the distribution of host trees. We also expect to find indicator species for each ecoregion.
ii. ECM fungi groups with long-distance dispersal (basidiomycetes with aboveground fruiting bodies and wind-dispersed spores) showing more homogenous communities (i.e., less marked distance decay and a weaker ecoregion effect) than hypogeous ascomycetes and basidiomycetes with resupinate fruiting bodies. We also expect a stronger ecoregion effect for communities associated with tree species showing a smaller latitudinal extent of distribution.
iii. a better resolution of biogeographic patterns when including species abundance, compared to the commonplace use of species occurrences (presence/absence) only.

## Materials and methods

### Studied region and plots description

We used a network of 129 forest plots, belonging to the intensively monitored, long-term Level II forest plot network of the UNECE International Co-operative Programme on Assessment and Monitoring of Air Pollution Effects on Forests (ICPF; http://icpforests.net; Ferretti, 2013), covering 12 contiguous ecoregions (as defined by Dinerstein et al., 2017) of Central and Northern Europe. Each forest plot was dominated by one of four host tree species: Scots pine (*Pinus sylvestris,* n = 41) and European beech (*Fagus sylvatica,* n = 30) both sampled in nine ecoregions, and Norway spruce (*Picea abies,* n = 36) and sessile or pedunculate oaks (*Quercus robur* and *Q. petrea, n* = 22) sampled in seven ecoregions (Table 1). Oak sites were pooled for the analyses as the distribution of the two species extensively overlap, they shared ca 64% of the taxa occurring more than once in the dataset and taxa present in at least five sites were detected in the roots of both oaks (Suz et al., 2014). Polygon data for the spatial delimitation of ecoregions were obtained from https://ecoregions2017.appspot.com (Dinerstein et al., 2017). The largest region is the Scandinavian and Russian Taiga with 2,170,289 km^2^ and the smallest is the English Lowlands beech forest with 45,769 km^2^. The number of sites sampled varies from three in the Cantabrian mixed forest and Celtic broadleaf forest to 38 in the Western European broadleaf forest. The average distance between plots within ecoregions is 384km (min = 0.3, max = 1,300km) and the average distance between plots between different ecoregions is 1,066km (min = 62.5, max = 3,273km). See Table 1 for sites and ecoregion descriptions.

**Table 1.** Area, total number of sites, number of sites dominated by each tree species in each ecoregion, number of root tips sampled, measured OTU richness, and estimated sample completeness for each ecoregion.

| *Ecoregion* | *Area (km2)* | *N sites* | *Beech* | *Pine* | *Spruce* | *Oak* | *N roots* | *Richness* | *Completeness* |
| --- | --- | --- | --- | --- | --- | --- | --- | --- | --- |
| *Alps conifer and mixed forests* | 149.87 | 11 | 3 | 2 | 6 | 0 | 1920 | 378 | 0.92 |
| *Baltic mixed forests* | 114.577 | 5 | 4 | 1 | 0 | 0 | 1024 | 147 | 0.94 |
| *Cantabrian mixed forests* | 96.266 | 3 | 2 | 0 | 0 | 1 | 415 | 164 | 0.77 |
| *Carpathian montane forests* | 125.335 | 5 | 1 | 0 | 4 | 0 | 1090 | 159 | 0.95 |
| *Celtic broadleaf forests* | 210.027 | 3 | 0 | 2 | 0 | 1 | 595 | 71 | 0.96 |
| *Central European mixed forests* | 733.979 | 10 | 0 | 7 | 1 | 2 | 1903 | 281 | 0.94 |
| *English Lowlands beech forests* | 45.769 | 4 | 1 | 1 | 0 | 2 | 648 | 127 | 0.92 |
| *European Atlantic mixed forests* | 386.557 | 21 | 5 | 9 | 0 | 7 | 3676 | 303 | 0.97 |
| *Pannonian mixed forests* | 307.716 | 5 | 1 | 0 | 1 | 3 | 1210 | 237 | 0.92 |
| *Sarmatic mixed forests* | 850.296 | 13 | 2 | 7 | 4 | 0 | 2451 | 339 | 0.94 |
| *Scandinavian and Russian taiga* | 2.170.289 | 11 | 0 | 6 | 5 | 0 | 2483 | 300 | 0.96 |
| *Western European broadleaf forests* | 290.094 | 38 | 11 | 6 | 15 | 6 | 6517 | 587 | 0.96 |

### Fungal communities

Relative abundance of ECM fungi was obtained from van der Linde et al. (2018). In each plot, 288 mycorrhizal root tips were collected along 20-24 transects (∼ 1 to 5 m depending on forest density) between two individuals of the dominant tree species. Individual mycorrhizas were cleaned, and the identity of the host tree was checked by visual inspection of the root morphology. From each individual mycorrhiza, DNA was extracted using Extract-N-AMP (Sigma) and the internal transcribed spacer (ITS) region of the rDNA was amplified using primers ITS1F and ITS4 (Gardes & Bruns, 1993; White et al., 1990). The amplicons were sequenced using the Sanger method. The obtained DNA sequences were assembled, filtered (>100bp) and matched to the UNITE database v 7.0, (Abarenkov et al., 2023) at a 97% similarity threshold. The unmatched sequences were clustered into *de novo* OTUs with the same threshold and assigned a taxonomic level based on their similarity with the UNITE reference database (see van der Linde et al. (2018) for details). In total, 23,932 mycorrhizas were assigned to 1,350 ECM fungal taxa. The median number of mycorrhizas identified per site was 195 (min = 55, max = 273). The dataset was composed of 1224 OTUs of Basidiomycota (19,509 mycorrhizas, 81.5%) and 126 OTUs of Ascomycota (4423 mycorrhizas, 18.5%). For each OTU, the fruitbody type was determined using expert knowledge and is provided in Supplementary material Table S1.

### Statistical analyses

All analyses were performed using the statistical computing environment R, version 4.3. (R Core Team, 2023). We estimated the completeness of the sampling for each ecoregion based on a species accumulation curve using the ‘iNEXT’ package (Hsieh et al., 2022).

To test if ECM fungal communities show a distance decay pattern, we modelled the similarity in composition as a negative exponential function of the geographic distance. We tested the model fit using a permutation method following Gómez-Rodríguez & Baselga (2018). The negative exponential function was chosen as it is the classical theoretical shape of distance decay at broad spatial scales encompassing large environmental variations (Nekola and McGill, 2014).

To test for differences in ECM fungal community composition among ecoregions, we used two complementary approaches. First, we built a dissimilarity matrix using the Morisita-Horn dissimilarity (function *vegdist*, package ‘*vegan’*, Oksanen et al., 2019) on the ECM fungal abundance data, which is robust towards under sampling (Jost et al., 2011), as is often the case with microbial data and allows the comparison of communities with different sample sizes. We then fit two models of distance decay for the pairs of plots belonging to the same ecoregion and for those belonging to different ecoregions. We quantified the strength of the ecoregion effect by comparing the intercepts of the curves of distance decays (calculated for the whole community) for pairs of plots within and between ecoregions, which is a way of measuring the difference in community similarity at a null geographical distance, using the *Zdep* statistic (Martín-Devasa et al., 2022). This method is analogous to a *t*-test on model parameters, based on a site-block permutation test of the observations (nperm = 1,000). Since the geographical distance between plots among ecoregions was usually larger than within ecoregions, we only kept the pairs of sites that are within the range of distance shared by both intra and inter pairs (between 62.5 and 1,300 km). To account for the differences in hosts between plots, we only kept the pairs of sites (intra- and inter-ecoregion) with the same dominant host tree. These two filtering steps resulted in 1,573 pairs of plots included in the analysis. To test for the importance of OTU abundance, we fit the same model on a distance decay matrix calculated using presence-absence data only. Similarly, we fit the same models on several subgroups of ECM fungi, to explore the effect of phylogeny, fruiting body type and host tree on ecoregion delimitation. P-values associated with the Zdep statistic of the different groups were adjusted for multiple tests using the Benjamini-Hochberg false discovery rate correction (Benjamini & Hochberg, 1995).

To test for pairwise differences in ECM community composition between ecoregions, we used a multivariate analysis of variance by permutation (PERMANOVA, function *adonis2*, ‘vegan’) using ecoregion as predictor followed by a *post hoc* test (function *pairwise.adonis2*, package ‘pairwiseAdonis’). P-values were adjusted using the Benjamini-Hochberg false discovery rate correction. To test if the results of the PERMANOVA were influenced by the dispersion of the data within ecoregion, we conducted a test of multivariate homogeneity of variance using the *betadisper* (‘vegan’) function followed by a Tukey test (*TukeyHSD* function). We graphically represented the differences between ecoregions using a non-metric multidimensional scaling (NMDS) ordination using the *metaMDS* function (package ‘vegan’). To investigate similarities between ecoregions, we also conducted a cluster analysis of ecoregions on a Morisita-Horn dissimilarity matrix of pooled ECM fungal communities (function *hclust*).

To disentangle the separate and joint effects of ecoregion, climate, and host tree, we conducted a variation partitioning on the distance matrix of community similarity (function *varpart*, package ‘vegan’). Mean annual temperature and precipitation are key determinants of ECM fungal communities (van der Linde et al., 2018) and, together with temperature and precipitation seasonality, explain differences in ecoregions (Smith et al., 2020). Mean annual value and seasonality for temperature (MAT, MST) and precipitation (MAP, MSP) of the 1970-2000 period were extracted for each site from the Worldclim 2 database (Fick and Hijmans, 2017) at a 30s spatial resolution. We used partial distance-based redundancy analysis (db-RDA) to quantify the fractions related to the different explanatory components of variation partitioning, using adjusted coefficients of determination and p-values obtained by permutation (Borcard et al., 2018).

Finally, as a complementary test of the ecological coherence of ecoregions, we tested for the presence of ecoregion-specific indicator species (*IndVa*l analysis) as well as species with high ‘specificity’ (A > 0.8) (Dufrêne and Legendre, 1997) using the *multipatt* function in the package ‘indicspecies’ with 50,000 permutations (De Cáceres et al., 2012). While species of high specificity are characterised as having most of their distribution in only one ecoregion, indicator species display both high specificity and high fidelity (Dufrêne and Legendre, 1997), making these two metrics ecologically complementary. Specificity and indicator species analyses were run based on OTUs present in at least two plots and with at least 10 mycorrhizas (277 OTUs). These analyses were conducted for the total dataset and by host tree. We only focus on *IndVal* values > 0.5 and Benjamini-Hockberg adjusted p-values to discuss the results.

## Results

### Distance decay of community similarity is driven by ecoregion and host species

The general shape of community distance decay across Europe using abundance data has a negative exponential shape with a low similarity at a zero distance (β_0_ = 0.21) and a moderate slope of decay of (β_dist_ = -0.40). When comparing only pairs of sites dominated by the same tree species, the intercept is higher and the slope steeper (β_0_ = 0.40, β_dist_ = -0.49, Table 2, Fig. S1a, b in Supporting information), suggesting a higher similarity at short distance and a more pronounced distance decay. Using occurrence data and controlling for tree species, the similarity in communities is lower and the distance decay is less clear (Fig. S1d).

**Table 2.** Parameters of the distance decay exponential curves for different taxonomic and functional categories of ectomycorrhizal fungi. Intercept and slope are the parameters of the general curve of distance decay for the group, intra is the intercept of the model for pairs of plots within the same ecoregion and inter is the intercept of the model for pairs of plots in different ecoregions. P-adj is the p-value adjusted for multiple comparisons using the Benjamini- Hochberg method.

|  | Group | Intercept | Slope | Intra | Inter | Zdep | p-adj |
| --- | --- | --- | --- | --- | --- | --- | --- |
| Total | Abundance | 0.40 | -0.49 | 0.34 | 0.51 | -4.00 | **<0.001** |
|  | Presence-absence | 0.30 | -0.44 | 0.28 | 0.34 | -3.36 | **0.002** |
| Fruiting body | Epigeous | 0.40 | -0.53 | 0.32 | 0.52 | -4.21 | **<0.001** |
|  | Hypogeous | 0.50 | -0.28 | 0.42 | 0.51 | -1.85 | 0.103 |
| Type | Mushroom | 0.42 | -0.80 | 0.35 | 0.54 | -3.64 | **0.001** |
|  | Sclerotium | 0.61 | 0.06 | 0.54 | 0.67 | -2.73 | **0.013** |
|  | Crust | 0.24 | -0.02 | 0.20 | 0.34 | -3.40 | **0.002** |
|  | Truffle | 0.52 | -0.50 | 0.45 | 0.44 | 0.20 | 0.984 |
| Phylum | Basidiomycota | 0.39 | -0.52 | 0.32 | 0.53 | -4.21 | **<0.001** |
|  | Ascomycota | 0.47 | -0.30 | 0.41 | 0.49 | -1.69 | 0.138 |
| Order | Russulales | 0.43 | -0.72 | 0.33 | 0.53 | -3.67 | **0.001** |
|  | Atheliales | 0.41 | -0.07 | 0.35 | 0.61 | -4.16 | **<0.001** |
|  | Telephorales | 0.23 | -0.54 | 0.20 | 0.28 | -1.92 | 0.096 |
|  | Agaricales | 0.23 | -0.59 | 0.22 | 0.21 | 0.11 | 0.993 |
|  | Boletales | 0.48 | -0.94 | 0.44 | 0.48 | -0.53 | 0.783 |
|  | Cantharellales | 0.18 | -0.51 | 0.16 | 0.28 | -1.18 | 0.333 |
|  | Pezizales | 0.36 | -0.69 | 0.41 | 0.36 | 0.48 | 0.784 |
| Host tree | *Pinus sylverstris* | 0.46 | -0.67 | 0.37 | 0.57 | -1.92 | 0.096 |
|  | *Picea abies* | 0.36 | -0.33 | 0.23 | 0.54 | -4.30 | **<0.001** |
|  | *Fagus sylvatica* | 0.34 | -0.31 | 0.33 | 0.31 | 0.07 | 0.993 |
|  | *Quercus* spp. | 0.48 | -0.58 | 0.80 | 0.81 | 0.00 | 1.000 |

Overall, there is a strong ecoregion effect, as illustrated by the large difference in intercepts of the distance decay curves within and between ecoregions (Fig. 2A). These results show that ECM fungal communities within the same ecoregion are 33.3% more similar to one another than they are to communities in a different ecoregion (Fig. 2A, Table 2). There is no clear difference in the slopes of both curves (Zdep = 1.95, p-adj > 0.05), suggesting that the rate of distance decay within and between ecoregions is similar. Using presence-absence data, there is also an ecoregion effect, although the overall similarity within and between ecoregion is smaller and the ecoregion effect accounts only for 17.6% of the community similarity (Fig. 2b, Table 2).

**Figure 1.**
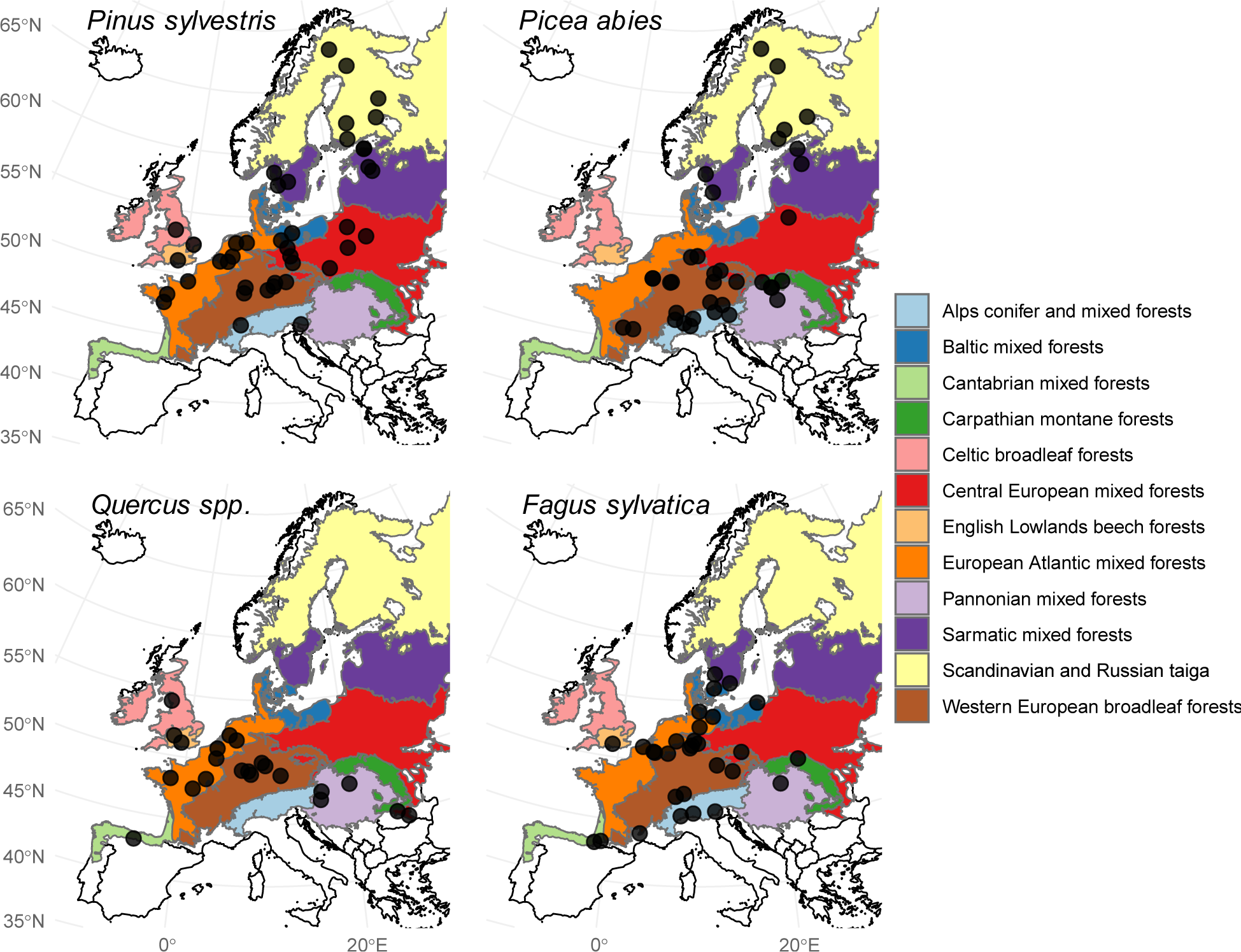
Delimitation of ecoregions and location of study plots per host tree species across Europe. *Pinus sylvestris* (n = 41), *Picea abies (*n = 36), *Quercus robur* & *Q. petraea (*n = 22) and *Fagus sylvatica* (n = 30).

**Figure 2.**
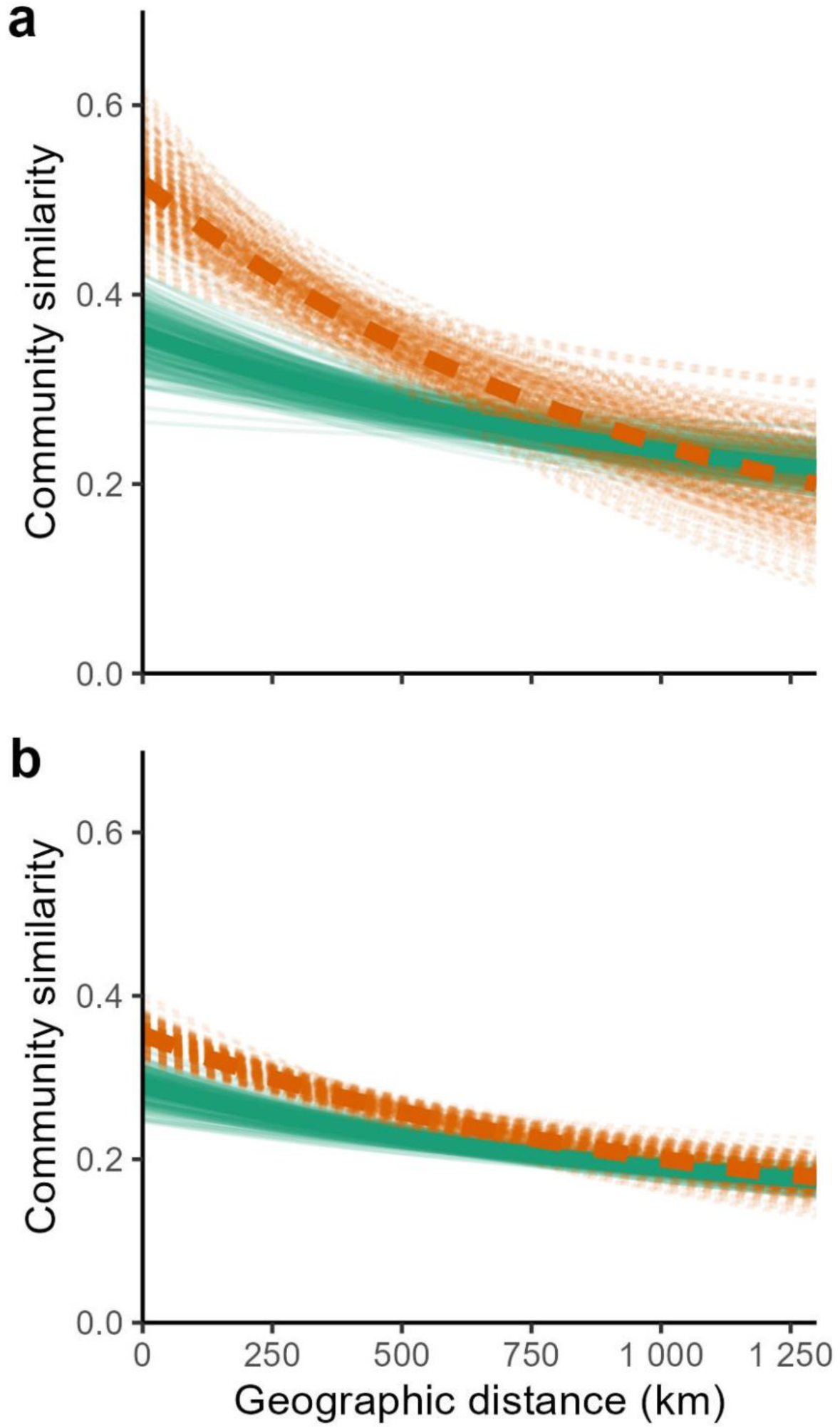
Distance decay curves of ectomycorrhizal fungal communities within (orange dashed line) and between (solid green line) ecoregions. The set of thin lines represents the incertitude calculated from 1,000 site-block permutations. The similarity was calculated including (a) or excluding (b) OTUs abundances (see Table 2 for curves parameters).

This method also highlights clear ecoregion effects for different subgroups of ECM fungi (Table 2). Fungi with aboveground fruiting bodies (epigeous) show a clear ecoregion effect, which is not the case for fungi with belowground fruiting bodies (hypogeous). This is partly explained by the fact that among the epigeous species, both mushrooms and crusts exhibit an ecoregion effect, while among the hypogeous ones, truffle-forming fungi do not show any ecoregional patterns. This is also because most epigeous fruiting bodies belong to the phylum Basidiomycota (Δ β_0_ = 0.21, p-adj < 0.001), while most of the hypogeous fungi belong to Ascomycota (Δβ_0_ = 0.07, p-adj = 0.138). Within Basidiomycota, only the ecologically dominant orders Russulales and Atheliales (corresponding to 51.8% and 33.0% of the OTUs in the dataset, respectively) show a strong ecoregion effect (Table 2).

Regarding the number of OTUs associated with the different host species, 541 OTUs are associated with *F. sylvatica*, 485 with *P. sylvestris*, 545 with *P. abies* and 496 with *Quercus spp*. When analysing the ECM fungi communities of the different host trees separately, *P. sylvestris* and *Quercus spp.* communities show more pronounced distance decay (β_dist_ = -0.67, R^2^ = 0.10, p-adj = 0.004 and β_dist_ = -0.58, R^2^ = 0.15, p-adj = 0.004, respectively) than *P. abies* and *F. sylvatica* (β_dist_ = -0.33, R^2^ = 0.07, p-adj = 0.004 and β_dist_ = -0.31, R^2^ = 0.02, p-adj = 0.004, respectively), but none of these differences are statistically significant (Table S2). There is a strong ecoregion effect for the ECM fungi communities associated with conifers, but only statistically significant at α < 0.1 (after Benjamini-Hochberg correction) for *P. sylvestris*. No effect is observed for communities associated with broadleaves (Table 2).

### The similarity between ECM fungal community composition varies between pairs of ecoregions

The PERMANOVA shows a significant effect of the ecoregion on the community composition (F_11, 117_ = 2.98, R^2^ = 0.22, p < 0.001). The pairwise PERMANOVA reveals complex patterns among ecoregions (Table 3, Fig S2). The Scandinavian and Russian taiga and European Atlantic mixed forests are the most distinct, being significantly different from six and five other ecoregions respectively, followed by the Western European broadleaf forests (different to three other ecoregions) and Sarmatic mixed forests, Pannonian mixed forests, Central European mixed forests and Alps conifer and mixed forests (different to two other ecoregions). Interestingly, the Baltic mixed forests, Cantabrian mixed forests, Celtic broadleaf forests and English Lowlands beech forests are not statistically significantly different from any other ecoregion. The largest differences in effect size are between the Scandinavian and Russian taiga and English Lowlands beech forests (R^2^ = 0.47), Cantabrian mixed forests (R^2^ = 0.44) and Celtic broadleaf forests (R^2^ = 0.40). The PERMDISP analysis shows that multivariate variance is not significantly different between ecoregions, except for the Scandinavian and Russian taiga, which has a significantly smaller variance in community composition than several other ecoregions (Table S3). The NMDS ordination representing the community dissimilarities between ecoregions is in supplementary material (Fig. S2). The cluster analysis shows four groups of ecoregions: 1) a western cluster composed of the European Atlantic mixed forests, English Lowlands beech forests, Baltic mixed, and Celtic broadleaf forests, 2) a northern cluster composed of the Scandinavian and Russian taiga and Sarmatic mixed forests, 3) a southern cluster including both Cantabrian and Pannonian mixed forests and, 4) a central cluster containing the Western European broadleaf forests , Carpathian montane forests, Central European mixed forests, and Alps conifer and mixed forests (Fig. 3).

**Figure 3.**
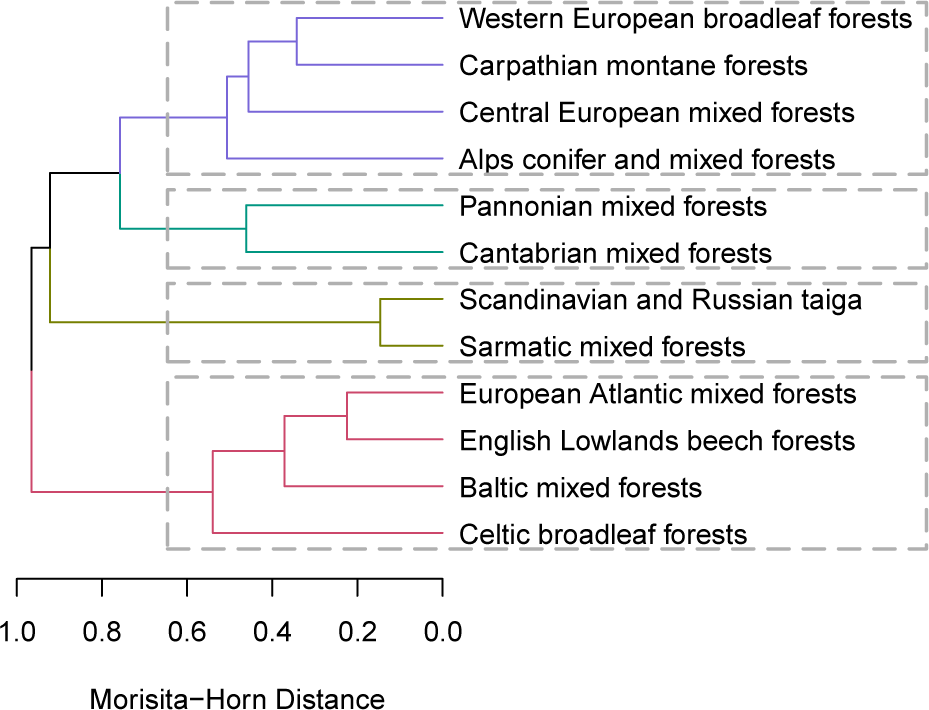
Hierarchical cluster analysis of the ectomycorrhizal composition of the 12 ecoregions studied, based on a Morisita-Horn dissimilarity matrix. The four groups discussed in the results are highlighted with boxes.

**Table 3.** Differences in ectomycorrhizal fungal community composition among ecoregions (pairwise PERMANOVA on Morisita-Horn dissimilarities). The lower triangle gives the R^2^ values, and the upper triangle represents the p-values adjusted for multiple comparisons using the Benjamini-Hochberg false discovery rate correction. The R^2^ of the pairs of ecoregions with p < 0.1 are bolded (p < 0.1., p<0.05*, p<0.01**).

|  | *Alps conifer*<br>*and mixed*<br>*forests* | *Baltic*<br>*mixed*<br>*forests* | *Cantabrian*<br>*mixed*<br>*forests* | *Carpathian*<br>*montane*<br>*forests* | *Celtic*<br>*broadleaf*<br>*forests* | *Central*<br>*European mixed*<br>*forests* | *English*<br>*Lowlands beech*<br>*forests* | *European*<br>*Atlantic mixed*<br>*forests* | *Pannonian*<br>*mixed*<br>*forests* | *Sarmatic*<br>*mixed*<br>*forests* | *Scandinavian*<br>*and Russian*<br>*taiga* | *Western*<br>*European*<br>*broadleaf forests* |
| --- | --- | --- | --- | --- | --- | --- | --- | --- | --- | --- | --- | --- |
| *Alps conifer and mixed forests* |  | 0.270 | 0.503 | 0.136 | 0.280 | 0.287 | 0.214 | 0.015 | 0.114 | 0.287 | 0.011 | 0.013 |
| *Baltic mixed forests* | 0.13 |  | 0.275 | 0.895 | 0.772 | 0.941 | 1.000 | 0.579 | 0.287 | 0.253 | 0.280 | 0.782 |
| *Cantabrian mixed forests* | 0.11 | 0.26 |  | 1.000 | 0.600 | 0.433 | 0.253 | 0.052 | 0.337 | 0.280 | 1.000 | 0.091 |
| *Carpathian montane forests* | 0.10 | 0.12 | 0.27 |  | 1.000 | 0.287 | 1.000 | 0.924 | 0.244 | 0.287 | 0.042 | 0.433 |
| *Celtic broadleaf forests* | 0.16 | 0.14 | 0.36 | 0.15 |  | 0.280 | 0.655 | 0.287 | 0.337 | 0.106 | 0.108 | 0.270 |
| *Central European mixed forests* | 0.08 | 0.10 | 0.15 | 0.10 | 0.10 |  | 0.098 | 0.026 | 0.136 | 0.075 | 0.007 | 0.280 |
| *English Lowlands beech forests* | 0.15 | 0.08 | 0.30 | 0.17 | 0.16 | 0.13. |  | 0.579 | 0.136 | 0.121 | 0.280 | 0.287 |
| *European Atlantic mixed forests* | 0.13* | 0.05 | 0.10. | 0.08 | 0.05 | 0.09* | 0.03 |  | 0.011 | 0.007 | 0.008 | 0.084 |
| *Pannonian mixed forests* | 0.12 | 0.24 | 0.17 | 0.22 | 0.27 | 0.13 | 0.26 | 0.13* |  | 0.280 | 0.280 | 0.007 |
| *Sarmatic mixed forests* | 0.07 | 0.15 | 0.18 | 0.12 | 0.16 | 0.09. | 0.19 | 0.16** | 0.18 |  | 0.015 | 0.106 |
| *Scandinavian and Russian taiga* | 0.24* | 0.37 | 0.44 | 0.37* | 0.40 | 0.22** | 0.47 | 0.31** | 0.41 | 0.10* |  | 0.007 |
| *Western European broadleaf forests* | 0.05* | 0.03 | 0.05. | 0.03 | 0.04 | 0.05 | 0.04 | 0.04. | 0.07** | 0.06 | 0.16** |  |

### The ecoregion effect is partly explained by climate but is independent of host tree

The variation partitioning of community similarity among climate, host and ecoregion explains a total of 36.6% of the pairwise similarity in ECM fungal community composition variation (Fig. 4). The total host tree fraction explains 22.6% of the variation (F_3, 125_ = 13.4, p < 0.001), followed by ecoregion explaining 14.7% (F_11, 117_ = 2.97, p < 0.001) and climate explaining 10.7% (F_4, 124_ = 4.83, p < 0.001). About half of the ecoregion variation is explained by the differences in climate while the host tree variation is mostly independent from ecoregion and climate (Fig. 4).

**Figure 4.**
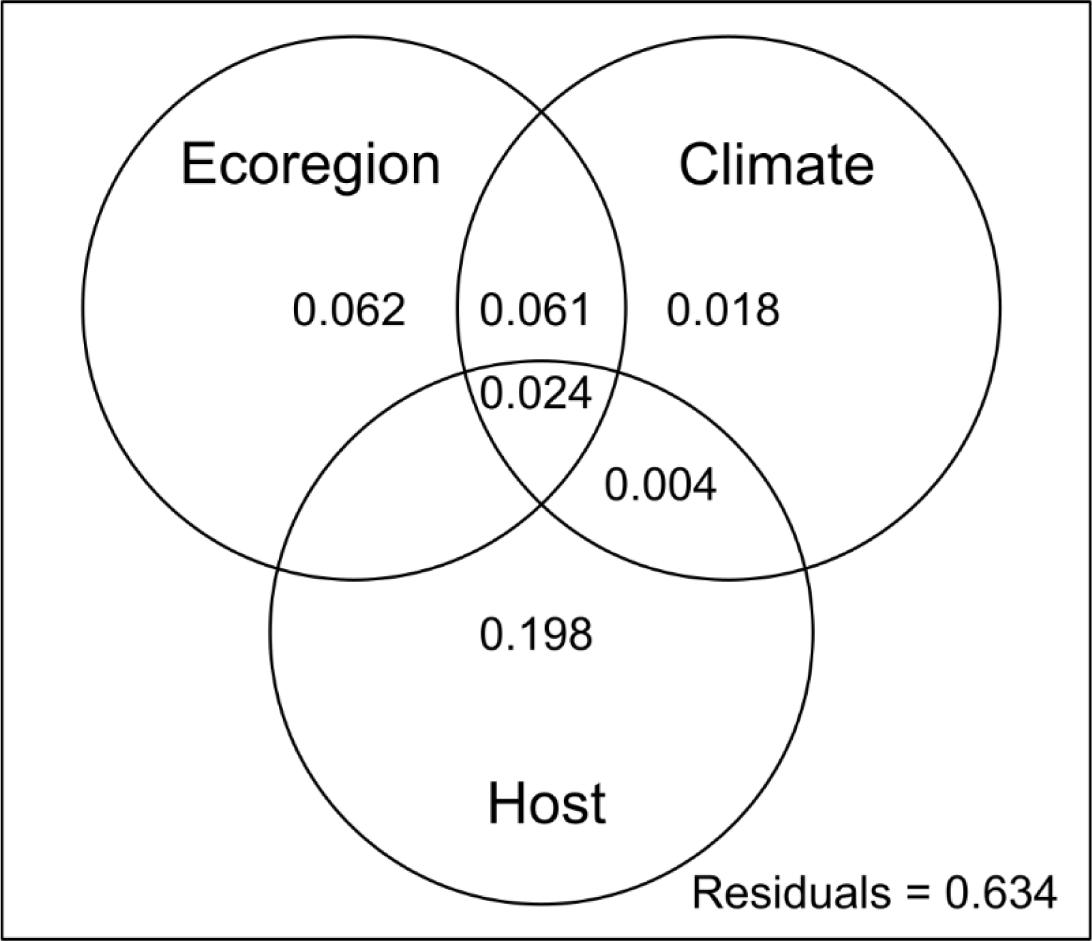
Variation partitioning of European ectomycorrhizal fungal communities between ecoregion, climate (mean annual value and seasonality of temperature and precipitation) and host tree identity. Values inside the circles are adjusted R^2^. Values < 0 are not shown.

### Ectomycorrhizal fungal species as indicators of ecoregions

Using the whole dataset, five OTUs emerge as statistically significant indicators (*IndVal* > 0.5, adjusted p-value < 0.05): two OTUs are indicator for the Scandinavian and Russian taiga and Pannonian mixed forests while one OTU is an indicator of the Cantabrian mixed forests. Looking at tree specific communities, three OTUs are indicators of the Scandinavian and Russian taiga, two are indicators of the Carpathian montane forests, and one is indicator of the European Atlantic mixed forests. At least one OTU shows a high specificity (A > 0.8) for each ecoregion (except the Western European broadleaf forests). The Scandinavian and Russian taiga and Pannonian mixed forests possess the highest number of specific OTUs with nine and seven respectively.

## Discussion

Biogeographical patterns are useful to study the eco-evolutionary processes, as well as to inform species conservation because they describe the outcome of complex interactions between biotic and abiotic components at large scales. Recent studies, have questioned the presence of large scale patterns in fungi, such as the distance decay in community similarity or the coherence of ecoregions, compared to plants of animals (Locey et al., 2020, Smith et al., 2020, 2018). These studies suggest that the discrepancies observed could result from the pooling of a the large diversity of organisms, potentially masking individual responses of different functional groups (Smith et al., 2018). However, a key limitation to study the biogeography of fungi is the lack of robust community data, resulting from the difficulties in quantifying fungal species abundance and distributions, particularly at large spatial scales. For example, non-standardised sampling (e.g. fruiting body records or eDNA), incomplete or inaccurate taxonomic identification, and indirect assignments of mycorrhizal status can all lead to errors and low-resolution data (Meyer et al., 2018; Suz et al., 2015).

Here, we found that ECM fungal communities are clearly structured by ecoregions and display a negative exponential distance decay in communities’ similarity at the European scale. Within ecoregions, communities are more similar than between ecoregions at similar geographical distances. This is partly explained by climate (ca. 57%), which suggest a role of environmental filtering, although other processes, such as dispersal limitation could be involved (Bowman & Arnolds, 2021). The large latitudinal extent of many ecoregions in our study could explain the weaker link between climate and ecoregions, compared to global studies (Tedersoo et al., 2022, 2014a). Further, other latitude-related variables, such as time since last glaciation, could also have remnant effects on geographical patterns in ECM fungi (Timling et al., 2012).

Discrepancies with previous studies (e.g. Smith et al., 2018, 2020) could be partly due to the inclusion in our study of a measure of functional abundance of the fungi; in fact, when calculating community similarity based on presence-absence, the ecoregion effect was weaker (Fig. 2). Including OTU abundance also increase the slope of distance decay in community similarity (Fig. S1b and d), as previously observed for North American ECM fungi (Bowman & Arnolds, 2021). Despite the large geographic distribution of some species, increasing similarity between communities, the change in their abundance and therefore their functional role in communities is key to understanding ecological as well as biogeographic patterns (Jost et al., 2011). This result is important because most recent microbial biogeographic studies (e.g. Vasar et al., 2022, Tedersoo et al., 2022) are based on soil metabarcoding (eDNA), where the relative abundance of each OTU cannot be readily compared among samples (Derocles et al., 2018 but see Shelton et al., 2023). This limitation in estimating effective abundance for ecosystem services, such as tree nutrition and carbon storage, will need to be more explicitly acknowledged and addressed in future research.

Our results also reveal different patterns of similarity between ecoregions, structured geographically, with some regions being more distinct and possessing indicator and specific OTUs. The Scandinavian and Russian taiga seems to be the most distinct in Europe, both by the high similarity of communities within and absolute differences in community composition with other ecoregions. This specificity can be explained both by geographic isolation due to the Baltic Sea, but also by the tree composition of these forests containing only conifer, compared to all other regions containing broadleaf trees.

### Traits and phylogeny influence ECM fungi biogeography

Within ECM fungi, the patterns in community composition of some more narrowly defined taxonomic and functional groups were also defined by ecoregions, with some exceptions (Table 2). For instance, the diversity of Ascomycota and the diversity of functional groups with a large share of Ascomycota, such as truffle-forming fungi, did not show a clear adherence to ecoregions, with generally higher similarity between communities and low coefficient of distance decay slopes. However, ECM communities in our dataset are composed of ca. 80% basidiomycetes and 20% ascomycetes, as observed previously in Europe and North America (Bowman & Arnold, 2021; Cox et al., 2010; Suz et al., 2014). This relatively low proportion of Ascomycota reduces the geographic resolution and statistical power of the analyses. The larger overall similarity and lower distance decay of ascomycetes and truffle-forming fungi could also be due to their dispersal agents (mostly animals, such as the Wild Boars), which can move their spores at a larger distance than for wind- or gravity-dispersed epigeous fungi, dispersing mostly at short distance (Johnson, 1996; Galante et al., 2011; Piattoni et al., 2016). Within basidiomycetes, the orders exhibiting a strong ecoregional pattern are Atheliales (*Piloderma* spp., *Amphinema* spp., *Tylospora* spp.) and Russulales (*Russula* spp., *Lactarius* spp.) representing 33.0% of the OTUs and 51.8% of all mycorrhizas in the dataset. The former produce inconspicuous crust-like fruiting bodies that are under-represented in global biodiversity databases (e.g. the Global Biodiversity Information Facility, GBIF). The fruiting bodies of these species, being small and adhering to the soil or the underside of woody debris, are likely to be inefficient for long distance dispersal, resulting in strong spatial patterns driven by dispersal limitation (Rosenthal et al., 2017). For Russulales, it has been shown that they could have a tropical origin (Hackel et al., 2022), which could result in many Russulales being more strongly structured along a climatic gradient and limited by low temperatures than other clades of ECM fungi that evolved mostly in temperate or boreal regions (Peay and Matheny, 2016). The absence of ecoregional structure for the other orders suggest, that species within these orders do not exhibit strong ecoregion preference. However, the proportion of species belonging to different orders can still vary among ecoregions because of their shared evolutionary history and niche conservatism. Therefore, investigating biogeographic patterns separately for groups of fungi exhibiting different dispersal traits and phylogenetic relationships is needed, as suggested previously for other microorganisms (Wetzel et al., 2012).

### Host species are central to biogeographic patterns in symbiotic organisms

Many biogeographic studies of fungi combine several host tree species. While this reflects the reality of natural ecosystems and the natural distribution of host trees in the environment, it can complicate the study of biogeographic patterns in species-rich communities of symbiotic organisms, as host identity is one of the main drivers of ECM fungi communities (Jarvis et al., 2013; van der Linde et al., 2018). We show that comparing only communities dominated by the same host species results in stronger biogeographical patterns (Fig. S1a and b). Communities associated with broadleaves trees show a weaker ecoregion effect compared to communities under conifers (Table 2). Because of a larger divergence between communities of different ecoregions, ECM fungi communities associated with conifer forests could be less resilient to environmental changes (Steidinger et al., 2020). However, this difference might also be due to the wider distributions of conifers compared to broadleaf trees in Europe and the higher susceptibility of ECM fungi that are conifer-specialists to certain environmental variables (van der Linde et al., 2018), leading to a tighter adherence to ecoregions by this group of fungi. Some difference in community composition, such as those observed between Scandinavian and Russian taiga and both Cantabrian and Pannonian mixed forests, could be partly explained by their differences in host tree compositions (Table 1) coupled with the specificity of a large proportion of ECM fungi to either broadleaves or conifers (van der Linde et al., 2018). But generally, this effect seems minor compared to other factors, such as distance and dispersal limitation, as suggested by the small (2%) joint effect of host and ecoregion in the variation partitioning.

### Limitations

Compared to studies of other groups of organisms such as insects (Kobayashi & Sota, 2016) or diatoms (Wetzel et al., 2012), the general similarity between all ECM fungi communities is low. This is likely due to a combination of factors. First, our study includes different host trees which harbour different ECM fungal communities, and only about half of ECM fungal species in our dataset are generalists (van der Linde et al., 2018). When comparing communities associated with the same host tree species, this similarity is much higher (Table 2). Second, the use of OTU as a proxy for species makes comparisons among kingdoms difficult. We used 97% similarity of the ITS region within OTUs, which is the most commonly used threshold but is not ideal for some clades (Wilson et al., 2023) such as *Cortinarius* (Garnica et al., 2016). This could explain the weak patterns observed for the Agaricales, within which *Cortinarius* represents a significant portion. This strict genetic similarity threshold is not easily compared with the species concepts applied in plants and animals; phylogenetic distance between species is variable, leading to inconsistent patterns of similarity between studies of micro- and macro-organismal ecology (Martin, 2002). This is the result of the difficulties to delimit single-locus molecular species in fungi to fit all purposes in ecological studies (Xu, 2020) – though this may need to be nuanced, as different similarity thresholds appear to have little influence on large-scale patterns of microbial species distribution (Botnen et al., 2018; Glassman et al., 2018). Finally, under sampling of some communities can also introduce a bias in comparing community similarity and diversity trends across space (Valdez et al., 2023). We estimate that the completeness of ECM fungal communities is sufficiently high to represent the abundant species well and we expect to have limited potential bias by using an abundance-sensitive measure of similarity (Jost et al., 2011). However, it is important to note that these results are influenced by the intensity of the sampling effort and the difference in sites numbers between ecoregions (Table 1). For example, the low explanatory power observed for the pairwise comparisons including the Western European broadleaf forest could partly arise from the large number of plots sampled in that region compared to other regions. This is commonly observed in large-scale studies (e.g. Tedersoo et al., 2022) which recently showed that endemicity (and therefore specificity) is less pronounced in Europe, where the sampling effort is greater than in other parts of the world.

## Conclusion

At the continental scale, climate, host, and soil characteristics are generally considered the main drivers of ECM fungal communities’ composition. Here, we found that ECM fungi follow a distance decay pattern and adhere to the ecoregion concept, independently from host and climate. This likely reflects key influences of both dispersal limitation and evolutionary processes. Major clades and functional groups of ECM fungi adhere to some ecoregion boundaries, although some ecoregions seem to be more distinct than others, which could reflect the influence of present and past climates. This new finding highlights the relevance of the widely recognised ecoregion concept for biogeographical analysis and incipient large-scale conservation planning for fungi. However, we suggest that including measures of the functional abundance of mycorrhizal fungi, and distribution of their hosts, are central to understand fungal biogeographic distribution and inform efficient conservation measures. Defining the most important factors that influence fungal distributions in Europe would require further investigation, to inform forest planning and aid conservation and climate change mitigation efforts, as well as defining hotspots regions for diversity and endemism. Future efforts need to focus on sampling ECM fungi globally, including a measure of functional abundance, using a standardised and robust procedure for integration of multiple datasets across fine to large scales. In addition, large scale genomic studies of key ECM fungal groups will allow the reconstruction of past dispersion and colonisation events to explain present-day biogeographic distributions and forecast their potential future.

## Acknowledgements

We acknowledge funding from NERC grant NE/ K006339/1 to M.I.B., from a Marie Skłodowska- Curie fellowship to LMS (FP7-PEOPLE-2009-IEF-253036), and from the European Union’s Horizon 2020 research and innovation programme under the Marie Skłodowska-Curie grant agreement no. 895799 to DB. We thank UNECE ICP Forests for access to forest plots. We thank Filipa Cox for providing mycorrhiza data for 12 plots. We thank D. Devey and L. Csiba for laboratory assistance; S. Boersma, F. van der Linde, H. van der Linde, J. van der Linde, C. Gonzales, A. Lenz, R. Lenz, S. Palacio, C. Palacio Castillo, S. Wipf, L. Garfoot, B. Spake, W. Rimington, J. Kowal, T. Solovieva, D. Gane, M. Terrington, J. Alden, A. Otway, V. Kemp, M. Edgar, Y. Lin, A. Drew, E. Booth, P. Cachera, R. De-Kayne, J. Downie, A. Tweedy, E. Moratto, E. Ek, P. Helminen, R. Lievonen, P. Närhi, A. Ryynänen, M. Rupel, J. Draing and F. Heun for field and laboratory work and R. Castilho for bioinformatics.

## Data availability

The sequences have been deposited on DDBJ/EMBL/GenBank under the BioProject PRJNA1066220, accession KIEV00000000. The unfiltered assembled sequences are available on Dryad at https://datadryad.org/stash/dataset/doi:10.5061/dryad.cr70qc8. The version described in this paper is the first version, KIEV01000000. The processed data and R code used for the analysis are available at https://github.com/guildelhaye/ecm_ecoregions. The GPS coordinates of each site is given at a geographic minute resolution following ICP Forest Publication Policy https://storage.ning.com/topology/rest/1.0/file/get/9870274098?profile=original

## Supporting information

Fig. S1. Distance decay curve of ectomycorrhizal communities in 129 sites across Europe encompassing 12 ecoregions.

Fig. S2. Non-metric multidimensional scaling representation of the ectomycorrhizal communities of the 12 studied ecoregions.

Table S1. Table of the taxonomy and traits for the 1,350 ectomycorrhizal fungal OTUs.

Table S2. Pairwise comparison of the distance decay parameters modelled by host tree species.

Table S3. Test of homogeneity of variance in ectomycorrhizal fungal community composition between ecoregions.

Table S4. Number of indicator and specific ectomycorrhizal fungal species in each ecoregion.

