## Supplementary material for "Ectomycorrhizal fungi are influenced by ecoregion boundaries across Europe"

The following Supporting Information is available for this article:

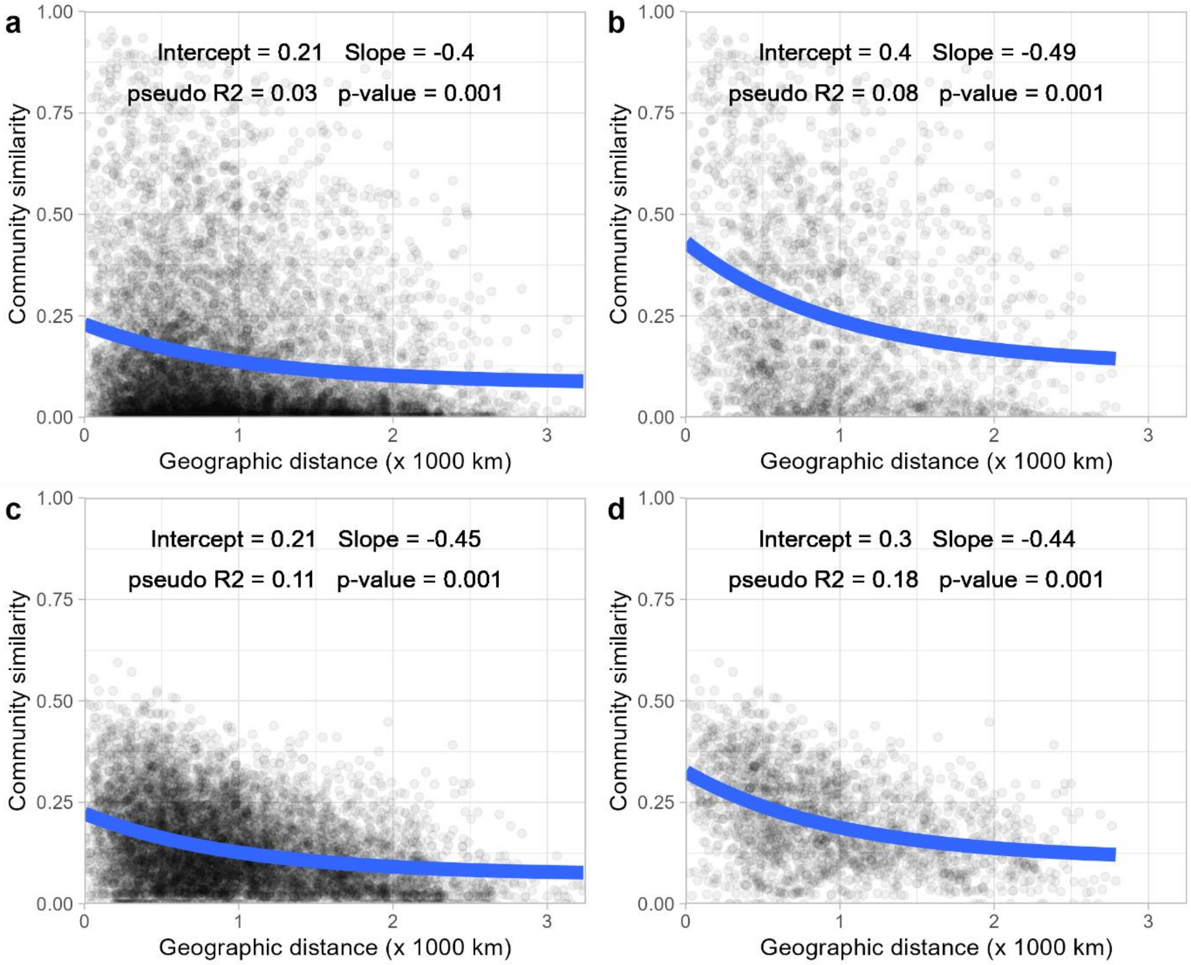


Fig. S1. Distance decay curve of the Morisita-Horn similarity in ectomycorrhizal communities in 129 sites across Europe encompassing 12 ecoregions. a,c) include all pairs of communities and b,d) include only the communities sharing the same host tree. a,b) abundance based dissimilarity and c,d) occurrence based dissimilarity.

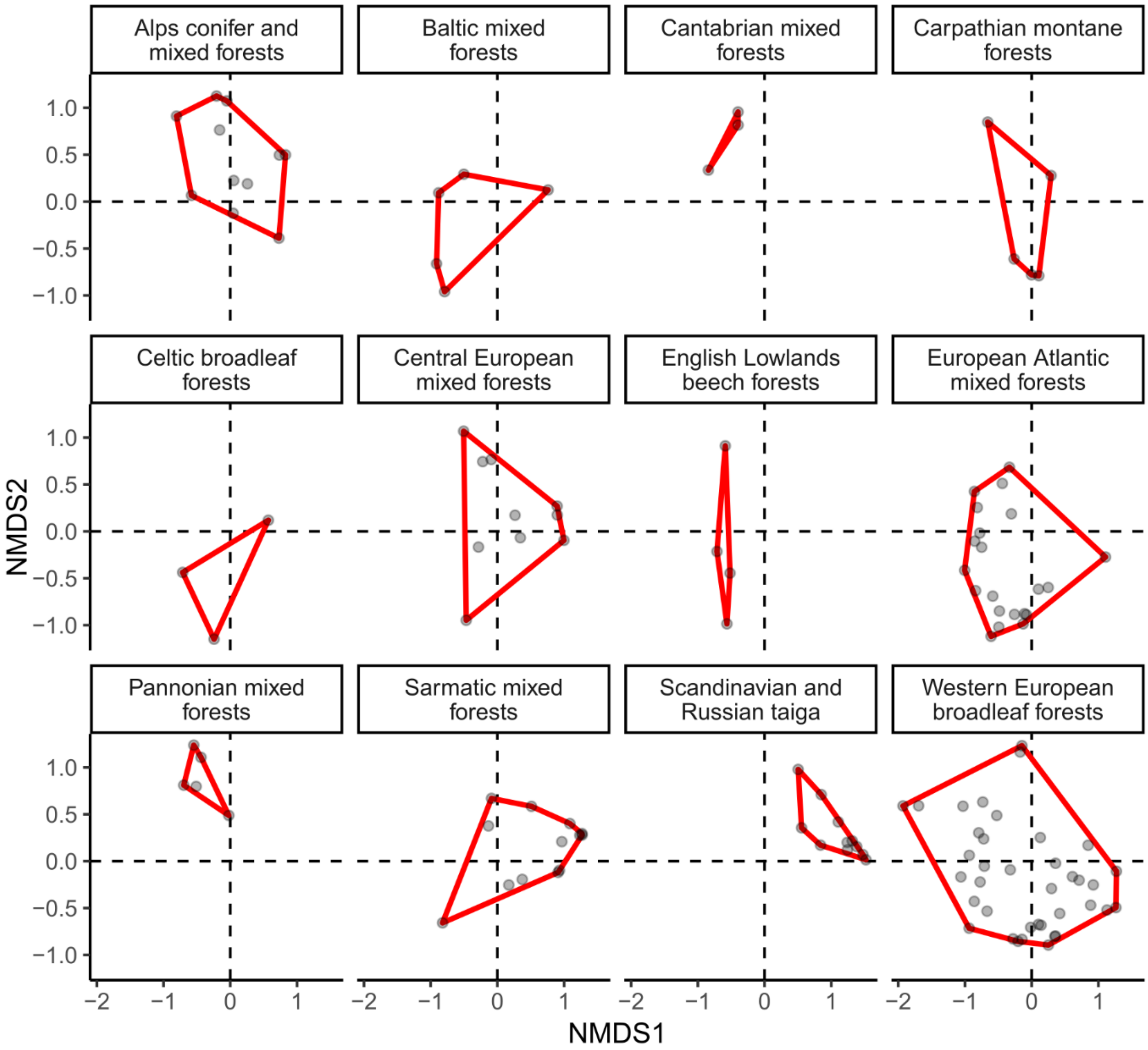


Fig. S2. Non-metric multidimensional scaling representation of the ectomycorrhizal communities of 12 ecoregions (stress = 0.09). Each dot represents a plot, and the red line represents the convex hull containing all plots of an ecoregion.

Table S1. Table of the taxonomy and traits for the 1,350 ectomycorrhizal fungal OTUs.

Table S2. Pairwise comparisons of model parameters between communities associated with different tree species. P-adj is the p-value adjusted for multiple comparisons using the Benjamini-Hochberg method.

| | trees | Tree 1 | Tree 2 | Zdep | p-adj |
|---|---|---|---|---|---|
| Intercept | oak_beech | 0.48 | 0.34 | 1.97 | 0.20 |
| | oak_pine | 0.48 | 0.46 | 0.27 | 0.78 |
| | oak_spruce | 0.48 | 0.36 | 1.76 | 0.20 |
| | beech_pine | 0.34 | 0.46 | -1.65 | 0.20 |
| | beech_spruce | 0.34 | 0.36 | -0.50 | 0.74 |
| | pine_spruce | 0.46 | 0.36 | 1.40 | 0.24 |
| Slope | oak_beech | -0.58 | -0.31 | -1.22 | 0.333 |
| | oak_pine | -0.58 | -0.67 | 0.39 | 0.840 |
| | oak_spruce | -0.58 | -0.33 | -1.33 | 0.333 |
| | beech_pine | -0.31 | -0.67 | 1.90 | 0.171 |
| | beech_spruce | -0.31 | -0.33 | 0.11 | 0.908 |
| | pine_spruce | -0.67 | -0.33 | -2.20 | 0.169 |

Table S3. Result of the Tukey test comparing the homogeneity of variance in community composition between each pair of ecoregions (PERMDISP).

| | diff | low | upp | adj_p |
|---|---|---|---|---|
| Baltic mixed forests-Alps conifer and mixed forests | -0.06 | -0.25 | 0.13 | 0.997 |
| Cantabrian mixed forests-Alps conifer and mixed forests | -0.16 | -0.40 | 0.07 | 0.458 |
| Carpathian montane forests-Alps conifer and mixed forests | -0.08 | -0.27 | 0.12 | 0.971 |
| Celtic broadleaf forests-Alps conifer and mixed forests | -0.05 | -0.29 | 0.18 | 0.999 |
| Central European mixed forests-Alps conifer and mixed forests | 0.04 | -0.11 | 0.20 | 0.998 |
| English Lowlands beech forests-Alps conifer and mixed forests | -0.15 | -0.36 | 0.06 | 0.387 |
| European Atlantic mixed forests-Alps conifer and mixed forests | -0.01 | -0.14 | 0.12 | 1 |
| Pannonian mixed forests-Alps conifer and mixed forests | -0.04 | -0.24 | 0.15 | 0.999 |
| Sarmatic mixed forests-Alps conifer and mixed forests | -0.04 | -0.18 | 0.11 | 0.999 |
| Scandinavian and Russian taiga-Alps conifer and mixed forests | -0.25 | -0.40 | -0.10 | 2.0E-05 |
| Western European broadleaf forests-Alps conifer and mixed forests | 0.03 | -0.09 | 0.15 | 0.999 |
| Cantabrian mixed forests-Baltic mixed forests | -0.11 | -0.37 | 0.16 | 0.972 |
| Carpathian montane forests-Baltic mixed forests | -0.02 | -0.25 | 0.21 | 1 |
| Celtic broadleaf forests-Baltic mixed forests | 0.01 | -0.26 | 0.27 | 1 |
| Central European mixed forests-Baltic mixed forests | 0.10 | -0.09 | 0.30 | 0.845 |
| English Lowlands beech forests-Baltic mixed forests | -0.09 | -0.34 | 0.15 | 0.975 |
| European Atlantic mixed forests-Baltic mixed forests | 0.05 | -0.13 | 0.23 | 0.998 |
| Pannonian mixed forests-Baltic mixed forests | 0.02 | -0.21 | 0.24 | 1 |

| | | | | |
|---|---|---|---|---|
| Sarmatic mixed forests-Baltic mixed forests | 0.02 | -0.17 | 0.21 | 0.999 |
| Scandinavian and Russian taiga-Baltic mixed forests | -0.19 | -0.38 | 0.00 | 0.058 |
| Western European broadleaf forests-Baltic mixed forests | 0.09 | -0.08 | 0.26 | 0.864 |
| Carpathian montane forests-Cantabrian mixed forests | 0.09 | -0.18 | 0.35 | 0.994 |
| Celtic broadleaf forests-Cantabrian mixed forests | 0.11 | -0.18 | 0.40 | 0.981 |
| Central European mixed forests-Cantabrian mixed forests | 0.21 | -0.03 | 0.44 | 0.143 |
| English Lowlands beech forests-Cantabrian mixed forests | 0.01 | -0.26 | 0.28 | 1 |
| European Atlantic mixed forests-Cantabrian mixed forests | 0.15 | -0.07 | 0.38 | 0.462 |
| Pannonian mixed forests-Cantabrian mixed forests | 0.12 | -0.14 | 0.38 | 0.921 |
| Sarmatic mixed forests-Cantabrian mixed forests | 0.13 | -0.10 | 0.36 | 0.793 |
| Scandinavian and Russian taiga-Cantabrian mixed forests | -0.09 | -0.32 | 0.15 | 0.986 |
| Western European broadleaf forests-Cantabrian mixed forests | 0.19 | -0.02 | 0.41 | 0.128 |
| Celtic broadleaf forests-Carpathian montane forests | 0.03 | -0.24 | 0.29 | 1 |
| Central European mixed forests-Carpathian montane forests | 0.12 | -0.07 | 0.32 | 0.644 |
| English Lowlands beech forests-Carpathian montane forests | -0.08 | -0.32 | 0.17 | 0.996 |
| European Atlantic mixed forests-Carpathian montane forests | 0.07 | -0.11 | 0.25 | 0.978 |
| Pannonian mixed forests-Carpathian montane forests | 0.04 | -0.19 | 0.26 | 0.999 |
| Sarmatic mixed forests-Carpathian montane forests | 0.04 | -0.15 | 0.23 | 0.999 |
| Scandinavian and Russian taiga-Carpathian montane forests | -0.17 | -0.36 | 0.02 | 0.138 |
| Western European broadleaf forests-Carpathian montane forests | 0.11 | -0.06 | 0.28 | 0.636 |
| Central European mixed forests-Celtic broadleaf forests | 0.10 | -0.14 | 0.33 | 0.968 |
| English Lowlands beech forests-Celtic broadleaf forests | -0.10 | -0.38 | 0.17 | 0.985 |
| European Atlantic mixed forests-Celtic broadleaf forests | 0.04 | -0.18 | 0.26 | 0.999 |
| Pannonian mixed forests-Celtic broadleaf forests | 0.01 | -0.25 | 0.27 | 1 |
| Sarmatic mixed forests-Celtic broadleaf forests | 0.02 | -0.21 | 0.25 | 1 |
| Scandinavian and Russian taiga-Celtic broadleaf forests | -0.20 | -0.43 | 0.04 | 0.191 |
| Western European broadleaf forests-Celtic broadleaf forests | 0.08 | -0.13 | 0.30 | 0.983 |
| English Lowlands beech forests-Central European mixed forests | -0.20 | -0.41 | 0.01 | 0.094 |
| European Atlantic mixed forests-Central European mixed forests | -0.05 | -0.19 | 0.08 | 0.979 |
| Pannonian mixed forests-Central European mixed forests | -0.09 | -0.28 | 0.11 | 0.950 |
| Sarmatic mixed forests-Central European mixed forests | -0.08 | -0.23 | 0.07 | 0.820 |
| Scandinavian and Russian taiga-Central European mixed forests | -0.29 | -0.45 | -0.14 | 4.9E-07 |
| Western European broadleaf forests-Central European mixed forests | -0.02 | -0.14 | 0.11 | 0.999 |
| European Atlantic mixed forests-English Lowlands beech forests | 0.14 | -0.05 | 0.34 | 0.375 |
| Pannonian mixed forests-English Lowlands beech forests | 0.11 | -0.13 | 0.35 | 0.923 |
| Sarmatic mixed forests-English Lowlands beech forests | 0.12 | -0.09 | 0.32 | 0.762 |
| Scandinavian and Russian taiga-English Lowlands beech forests | -0.10 | -0.31 | 0.11 | 0.931 |
| Western European broadleaf forests-English Lowlands beech forests | 0.18 | -0.01 | 0.37 | 0.069 |
| Pannonian mixed forests-European Atlantic mixed forests | -0.03 | -0.21 | 0.15 | 0.999 |
| Sarmatic mixed forests-European Atlantic mixed forests | -0.03 | -0.15 | 0.10 | 0.999 |
| Scandinavian and Russian taiga-European Atlantic mixed forests | -0.24 | -0.37 | -0.11 | 1.5E-06 |
| Western European broadleaf forests-European Atlantic mixed forests | 0.04 | -0.06 | 0.14 | 0.979 |
| Sarmatic mixed forests-Pannonian mixed forests | 0.00 | -0.18 | 0.19 | 1 |
| Scandinavian and Russian taiga-Pannonian mixed forests | -0.21 | -0.40 | -0.01 | 0.024 |
| Western European broadleaf forests-Pannonian mixed forests | 0.07 | -0.10 | 0.24 | 0.967 |
| Scandinavian and Russian taiga-Sarmatic mixed forests | -0.21 | -0.36 | -0.06 | 2.0E-4 |

| | | | | |
|---|---|---|---|---|
| Western European broadleaf forests-Sarmatic mixed forests | 0.07 | -0.05 | 0.18 | 0.759 |
| Western European broadleaf forests-Scandinavian and Russian taiga | 0.28 | 0.15 | 0.40 | 1.0E-09 |

Table S4. Number of OTUs with IndVal > 0.5; p-adj<0.05, (IndVal >0.5; p-adj<0.1) and [specificity > 0.8] for each tree species (*Pinus sylvestris*, *Picea abies*, *Fagus sylvatica*, *Quercus spp*.). Empty cells represent ecoregions with less than 2 plots of the tree considered.

| *Ecoregion* | All | *P. sylvestris* | *P. abies* | *F. sylvatica* | *Q. spp.* |
|---|---|---|---|---|---|
| *Alps conifer and mixed forests* | 0 (0) [1] | 0 (0) [1] | 0 (1) [3] | 0 (0) [2] | |
| *Baltic mixed forests* | 0 (0) [2] | | | 0 (0) [1] | |
| *Cantabrian mixed forests* | 1 (2) [1] | | | 0 (0) [1] | |
| *Carpathian montane forests* | 0 (1) [3] | | 2 (5) [3] | | |
| *Celtic broadleaf forests* | 0 (1) [4] | 0 (1) [2] | | | |
| *Central European mixed forests* | 0 (0) [2] | 0 (0) [6] | | | 0 (0) [0] |
| *English Lowlands beech forests* | 0 (0) [1] | | | | 0 (0) [1] |
| *European Atlantic mixed forests* | 0 (0) [2] | 1 (1) [4] | | 0 (0) [1] | 0 (0) [5] |
| *Pannonian mixed forests* | 2 (4) [7] | | | | 0 (2) [3] |
| *Sarmatic mixed forests* | 0 (0) [3] | 0 (0) [3] | 0 (0) [2] | 0 (0) [0] | |
| *Scandinavian and Russian taiga* | 2 (2) [9] | 2 (2) [7] | 1 (6) [4] | | |
| *Western European broadleaf forests* | 0 (0) [0] | 0 (0) [1] | 0 (0) [0] | 0 (0) [2] | 0 (0) [1] |
